# Super LeArner Prediction of NAb Panels (SLAPNAP): A Containerized Tool for Predicting Combination Monoclonal Broadly Neutralizing Antibody Sensitivity

**DOI:** 10.1101/2020.06.23.167718

**Authors:** D Benkeser, BD Williamson, CA Magaret, S Nizam, PB Gilbert

**Affiliations:** Department of Biostatistics and Bioinformatics, Emory University, Atlanta, GA 30322, USA; Vaccine and Infectious Disease Division, Fred Hutchinson Cancer Research Center, Seattle, WA 98109, USA; Vaccine and Infectious Disease and Public Health Sciences Divisions, Fred Hutchinson Cancer Research Center, Seattle, WA 98109, USA; Department of Biostatistics, University of Washington, Seattle, WA 98195, USA

**Keywords:** Broadly neutralizing antibodies, HIV-1, sieve analysis, efficacy trials, superlearning, variable importance estimation, proteomic antibody resistance score, ranking and down-selection, combination regimen, multi-specific regimen

## Abstract

**Summary:** Single broadly neutralizing antibody (bnAb) regimens are currently being evaluated in randomized trials for prevention efficacy against HIV-1 infection. Subsequent trials will evaluate combination bnAb regimens (e.g., cocktails, multi-specific antibodies), which demonstrate higher potency and breadth in vitro compared to single bnAbs. Given the large number of potential regimens in the research pipeline, methods for down-selecting these regimens into efficacy trials are of great interest. To aid the down-selection process, we developed Super LeArner Prediction of NAb Panels (SLAPNAP), a software tool for training and evaluating machine learning models that predict in vitro neutralization resistance of HIV Envelope pseudoviruses to a given single or combination bnAb regimen, based on Envelope amino acid sequence features. SLAPNAP also provides measures of variable importance of sequence features. These results can rank bnAb regimens by their potential prevention efficacy and aid assessments of how prevention efficacy depends on sequence features.

**Availability and Implementation:** SLAPNAP is a freely available docker image that can be downloaded from DockerHub (https://hub.docker.com/r/slapnap/slapnap). Source code and documentation are available at GitHub (respectively, https://github.com/benkeser/slapnap and https://benkeser.github.io/slapnap/).

**Contact:** David Benkeser

---

Extensive research has been conducted on passive administration of monoclonal broadly neutralizing antibodies (bnAb) with the objective of preventing HIV-1 infection (Morris and Mkhize, 2017). BnAbs target conserved epitopes on the HIV-1 envelope (Env) glycoprotein, and recent work has shown that combination bnAb regimens can neutralize most clinical HIV-1 isolates in genetically diverse Env panels [e.g., (McCoy and Burton, 2017; Sok and Burton, 2018)]. BnAb regimens have also been shown to prevent SHIV infection in nonhuman primate models (Gautam *et al.*, 2016; Hessell *et al.*, 2018; Pegu *et al.*, 2019). These developments position bnAbs as promising tools for the prevention of HIV-1, as well as other infectious diseases, in the near future (Karuna and Corey, 2020).

A key issue in developing efficacious bnAb regimens is understanding the neutralization breadth and potency of a given bnAb regimen against HIV-1 Env panels that are representative of circulating virus populations (Wagh *et al.*, 2016; Wagh *et al.*, 2018). Several bnAb combinations targeting distinct Env epitopes have been identified that exhibit greater neutralization breadth and potency than their constituent single bnAbs, with in vitro neutralization coverage rates approaching 100% (Doria-Rose *et al.*, 2012; Kong *et al.*, 2015; Wagh *et al.*, 2016). Thus, while the trials that are furthest advanced in the clinical pipeline [the Antibody Mediated Prevention trials (Gilbert *et al.*, 2017)] are evaluating the prevention efficacy of a single passively administered bnAb, VRC01, future trials will likely focus on bnAb combination regimens or multi-specific bnAb regimens. Several such early-phase clinical trials are being planned or are underway (e.g., NCT04212091, NCT03928821).

Analysis of efficacy trials can elucidate in vivo impact of neutralization breadth and sensitivity on prevention efficacy. Such analyses may lead to the validation of a bnAb-based surrogate endpoint for HIV-1 infection, which could accelerate the development of new prevention modalities, such as new bnAb regimens or novel vaccines that induce bnAbs (Liao *et al.*, 2013; Moody *et al.*, 2016; Williams *et al.*, 2017; Zhang *et al.*, 2016). To realize this exciting potential, analyses of randomized trials will need to be informed by in vitro analyses of bnAb breadth and potency. For example, in preparation for the sieve analysis of the AMP trials, Magaret et al. developed models predicting VRC01 neutralization sensitivity using Env amino acid (AA) sequence features (Magaret *et al.*, 2019), based on HIV-1 gp160 pseudoviruses from the Compile, Analyze and Tally NAb Panels (CATNAP) database (Yoon *et al.*, 2015). The super learning ensemble machine learning approach (van der Laan *et al.*, 2007) used to predict right-censored 50% inhibitory concentration titer (IC_50_) for each pseudovirus yielded a cross-validated area under the ROC curve (AUC) of 0.868. Magaret et al. also identified important sequence features for predicting VRC01 sensitivity with the goal to enable the AMP sieve analysis to focus on top-ranked features, thereby improving statistical power. Recently, Bricault et al. conducted similar variable importance signature analyses that generalized to all bnAbs across four antibody classes (Bricault *et al.*, 2019).

Given the movement towards combination or multi-specific bnAb regimens, we developed **Super LeArner Prediction of NAb Panels** (SLAPNAP), a publicly available, containerized pipeline that can perform an end-to-end analysis of in vitro neutralization data for bnAb combinations. SLAPNAP analyses of in vitro neutralization data can be used to inform the down-selection of combination or multi-specific bnAb regimens for future efficacy trials and in vivo analyses. SLAPNAP leverages all data available in the CATNAP database, and given a user-selected bnAb combination and neutralization endpoint, performs a suite of machine learning-based analyses and provides a report that summarizes the predictive results and highlights important pseudovirus sequence features.

## Methods

SLAPNAP is based on in vitro neutralization data available in the CATNAP database. Predictive analysis of neutralization sensitivity can be done for any single bnAb available in this database, or for combinations of any number of available bnAbs. In the latter case, the neutralization sensitivity of a pseudovirus to the combination bnAb regimen is estimated based on a Bliss-Hill model (Wagh *et al.*, 2016). The models created by SLAPNAP will predict one of three continuous measures of sensitivity [estimated IC_50_, estimated IC_80_, and instantaneous inhibitory potential (IIP) (Shen *et al.*, 2008)] and/or two binary measures of sensitivity [whether the estimated IC_50_ < a user-specified cutpoint, and whether the estimated IC_50_ < the cutpoint for a user-specified number of bnAbs in the specified combination (multiple sensitivity)]. Predictive models can be built using random forests (Breiman, 2001), boosted regression trees (Friedman, 2001), elastic net (Zou and Hastie, 2005), and/or a super learner ensemble (van der Laan *et al.*, 2007). Cross-validated tuning parameter selection is also available. Feature importance is estimated using the methods of Williamson et al. (Williamson *et al.*, 2020a; Williamson *et al.*, 2020b) and algorithm-specific importance measures (e.g., coefficient magnitude for the elastic net) can also be reported.

## Usage

SLAPNAP is a Docker container hosted on DockerHub (Docker Inc., 2019). With Docker installed, SLAPNAP can be downloaded by executing the following at the command line:

~~~
docker pull slapnap/slapnap:latest
~~~

SLAPNAP is executed using the `docker` run command. For example, the following code will instruct SLAPNAP to create and evaluate a neutralization predictor for the bnAb combination VRC07-523-LS and PGT121:

~~~
docker run \
  -v path/to/local/save/directory:/home/output/ \
  -e nab=“VRC07-523-LS;PGT121” \
  -e outcomes=“IC_50_;estsens” \
  -e learners=“rf;lasso” \
  -e importance_grp=“marg” \
  -e importance_ind=“pred” \
  slapnap/slapnap:latest
~~~

The `-v` tag specifies the directory on the user’s computer where the report will be saved, and `path/to/local/save/directory` should be replaced with the desired target directory. Options for the analysis are passed to the container via the `-e` tag; these options include the bnAbs to include in the analysis (`nab`), the neutralization outcomes of interest (`outcomes`), the learners to use in the analysis (`learners`), and the types of variable importance to compute (`importance_grp`, for groups of variables; `importance_ind`, for individual variables). Other output (e.g., the formatted analysis dataset and the fitted learners) can be requested via the `return` option. A full list of options and their syntax are available in the SLAPNAP documentation (https://benkeser.github.io/slapnap/).

Following the completion of a SLAPNAP run, the HTML report — along with any other requested output — will be saved to the output directory and the SLAPNAP container will shut down. Excerpts from the report are shown in Figure 1. Each report begins with a section summarizing data extraction and key results (Panel A). Descriptive statistics and plots are provided for each requested bnAb and for the estimated sensitivity measures for the combined bnAbs (Panel B). The performance of the learners for predicting binary sensitivity endpoints is estimated using cross-validated ROC analysis (Panel C). Feature importance is determined by a variety of methods and summarized in tables and figures (Panel D).

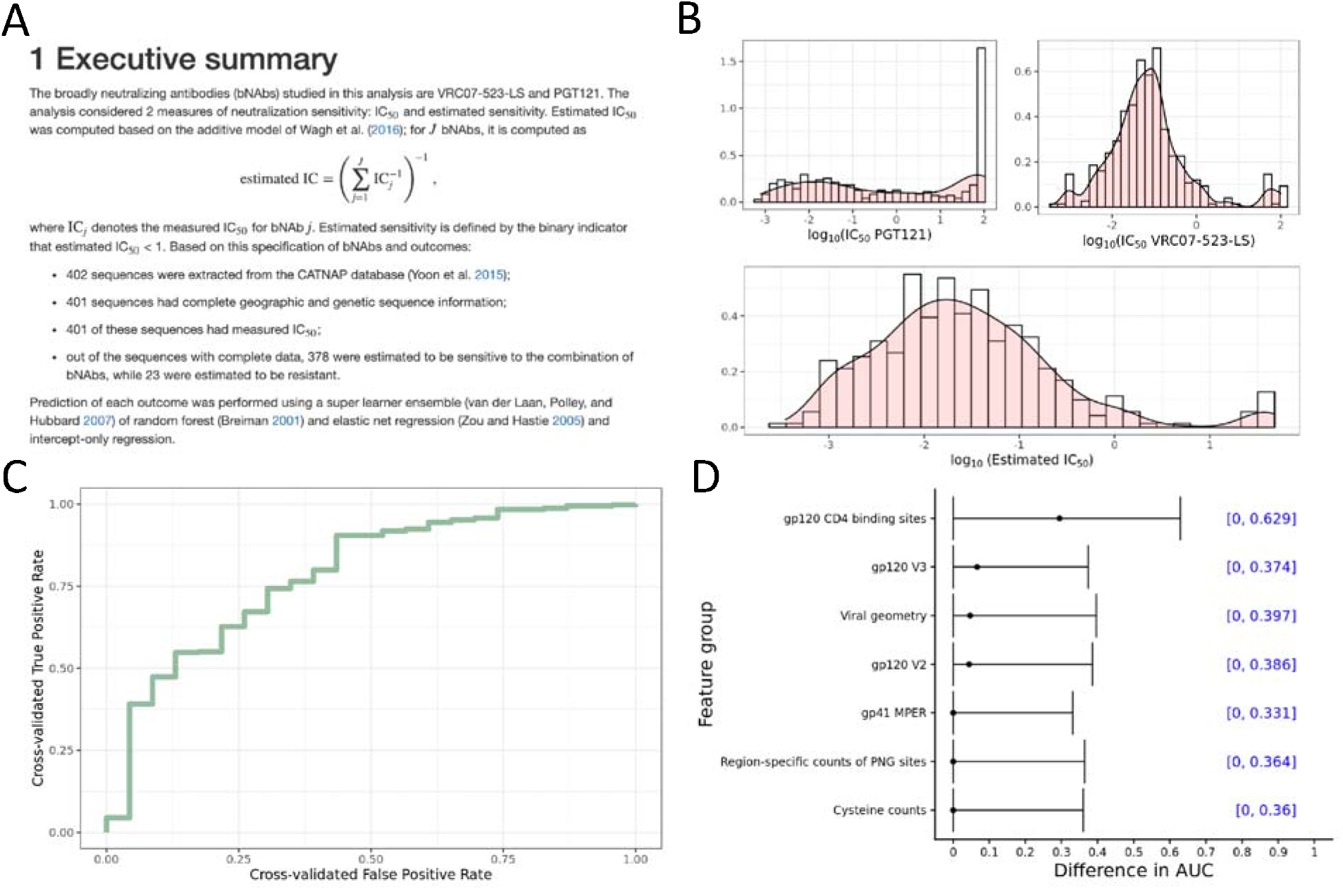

## Acknowledgements

**Conflict of interest:** None declared.

